# More than a feeling: central amygdala mediates social transfer of information about proximity of danger

**DOI:** 10.1101/2020.02.28.969113

**Authors:** K. Andraka, K. Kondrakiewicz, K. Rojek-Sito, K. Ziegart-Sadowska, K. Meyza, T. Nikolaev, A. Hamed, M. Kursa, M. Wójcik, K. Danielewski, M. Wiatrowska, E. Kublik, M. Bekisz, T. Lebitko, D. Duque, T. Jaworski, H. Madej, W. Konopka, P. M. Boguszewski, E. Knapska

## Abstract

To survive, an animal must adequately respond to challenges presented by the environment. Social animals can learn about danger from fear of conspecifics which allows them to avoid predation and other threats without costly, first-hand experience. However, it remains unclear as to whether animals can transmit specific information that helps an individual avoid harm or whether the transmitted social signals simply increase vigilance in a non-specific manner. Here we demonstrate that animals can select appropriate defensive strategies in a novel environment depending on cues from the conspecific and, using opsins targeted to behaviorally activated neurons, identify distinct neuronal circuits in the central amygdala (CeA) that are crucial for choosing a context-appropriate reaction. The identified circuits differ in molecular markers and patterns of connectivity. Thus, we show that social signals carry explicit information about proximity of danger, necessary for choosing a context-appropriate reaction, and that the choosing process is mediated by the CeA circuits.

## Introduction

When exposed to danger, animals display different behavioral strategies depending on the proximity of the threat. For example, detecting cues which potentially signal the presence of a predator usually triggers risk assessment behaviors such as rearing or brief episodes of exploration followed by runs^1^. Direct detection of a predator, on the other hand, typically evokes flight or freezing – depending on whether an escape route is available or not^1^. There is now a substantial body of literature describing which neural circuits mediate selection of an appropriate defensive reaction in individually tested animals^2,3^.

What is not known is how such decisions are influenced by social context. In group-living species, information about predators is often obtained through observation of conspecifics^4–6^. Under laboratory conditions this can be shown by exposing a non-stressed animal (called the ‘observer’) to a partner displaying freezing (a fearful ‘demonstrator’). This typically triggers freezing in the observer even in absence of any other aversive stimuli – a phenomenon known as ‘fear contagion’ or ‘observational fear’^4,7–10^. Such behavior is often interpreted as automatic and evolutionary hard-wired mimicry^7–9^ (but see also:^11^). Within this experimental framework it is difficult to determine whether warning signals emitted by conspecifics trigger only stereotypical defensive reactions, or if they can be used to derive information about proximity of danger and flexibly adjust the behavioral strategy to environmental challenges.

To test this, we compared two well established ‘fear contagion’ models in which a demonstrator provides information either about an imminent or a remote threat. Aside from studying behavioral reactions, we also investigated which central amygdala (CeA) circuits are activated in observers in each paradigm. We focused on this brain structure because of its well-established role in mediating single-subjects’ decisions about appropriate defensive reactions, such as freezing or flight^2,3,12^. As the CeA neuronal circuits critical for rapid selection of defensive responses have already been identified, we were able to investigate whether fear contagion shares neuronal circuits used for directly experienced fear. Using a combination of molecular, anatomical, and optogenetic techniques we characterized circuits implicated in each type of social behavior on the level of connectivity, molecular markers, and function.

## Results

### Social cues indicating imminent vs. remote danger trigger distinct behavioral responses

As a test, we used two behavioral models in which rats interacted with fearful cage mates. The fear they observed provided them with information about either imminent or remote danger. In the imminent threat paradigm, rats (called ‘observers’) witnessed their partners (‘demonstrators’) receiving electric foot shocks. In the remote threat paradigm, observers were exposed to demonstrators that had recently received foot shocks in an adjacent, sound-attenuated^13,14^ cage (Fig. 1A). Animals in both the imminent and remote threat models spent more time in social contact as compared to rats exposed to non-fearful partners (Fig. 1B,C; imminent threat: t=2.601 df=18, p=0.0180, remote threat: t=2.501 df=16, p=0.0236). The presence of the emotionally aroused partner in the imminent model significantly decreased exploratory behaviors such as locomotion and rearing (Fig. 1D,E; locomotion: U=10, p=0.0015; rearing: t=3.532, df=18, p=0.0024) while increasing defensive freezing (Fig. 1F; t=2.958, df=18, p=0.0084). The imminent paradigm also induced ultrasonic vocalizations (USVs) in the 22 kHz band, considered to be alarm calls (Fig. 1G,H; number: U=12, p=0.0098; frequency: U=0, p<0.0001). In contrast, interaction with an emotionally aroused partner in the remote threat model led to a significant increase in rearing (Fig. 1B,D; t=2.491 df=16, p= 0.0241). It also evoked high frequency ultrasonic vocalizations (50-70 kHz, Fig. 1G,H; number: t=2.406, df=7, p=0.0470) typically observed during peaceful social interactions^5^. Since freezing is generally performed when an animal recognizes danger as imminent^6^ and rearing is performed for information-gathering and risk assessment in response to environmental change^13^, the responses of observers in both models seem well suited to distance from threat and the context experienced by demonstrators.

**Fig. 1.**
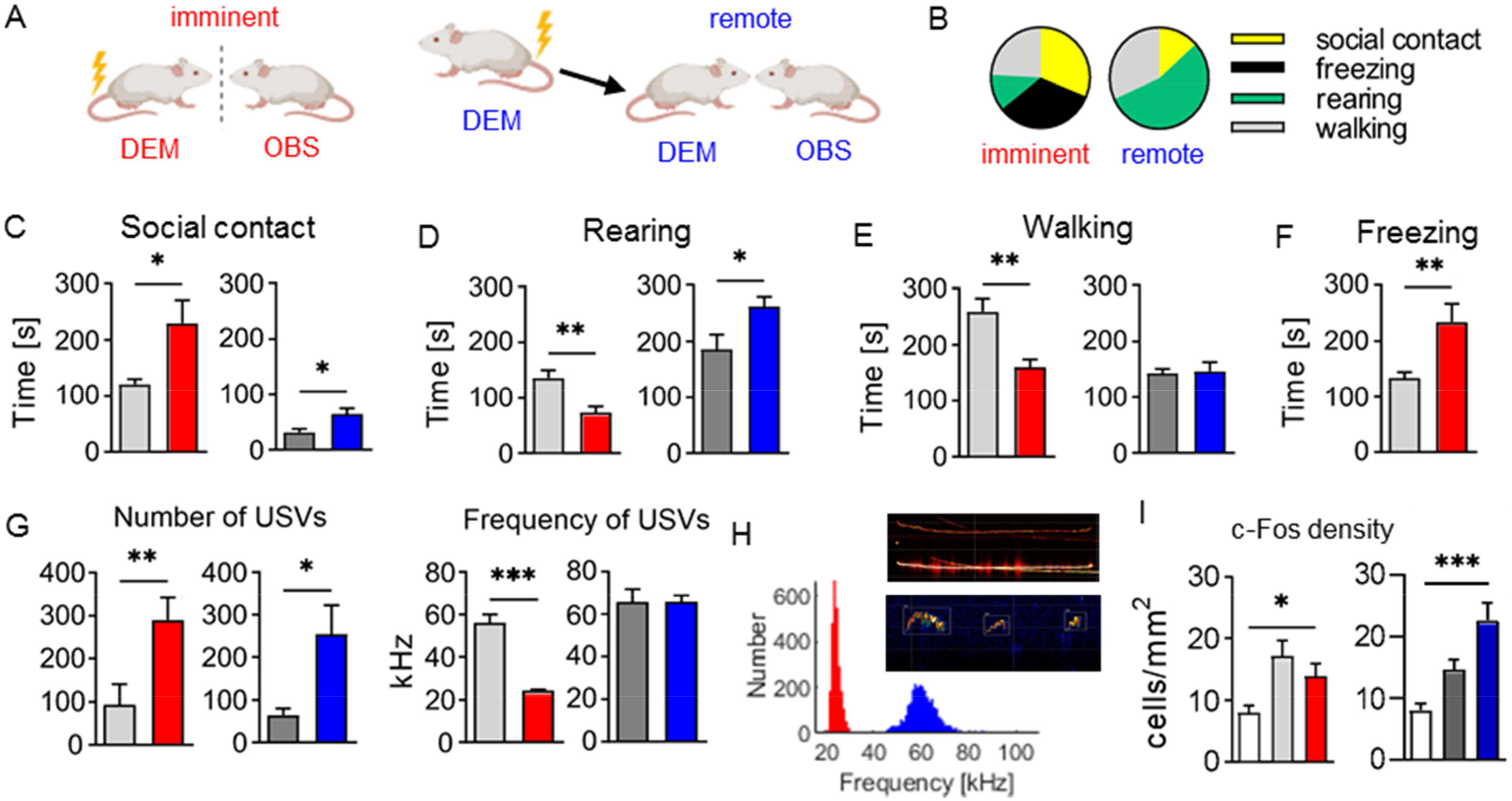
Rats respond differently to social information about imminent vs. remote threats. (**A**) Schematic representation of imminent and remote threat paradigms. (**B**) Observer rats manifest different patterns of behavior during imminent and remote paradigms - dominated by freezing and rearing, respectively. (**C**) In both models (red: imminent, blue: remote) observer rats spend more time on social contact than the control group (with non-stimulated partner; in gray). (**D**) Rearing is decreased in the imminent, but increased in the remote paradigm. (**E**) Walking decreases in the imminent paradigm. (**F**) Freezing increases in the imminent paradigm. Freezing is not observed in the remote paradigm. (**G**) Left panel: in both models, rats emit more ultrasonic vocalizations (USV) than control groups. Right panel: in the imminent paradigm, mean frequency of USV is decreased, while in the remote paradigm it is unchanged. (**H**) Vocalizations emitted in the two models are very different – long 22kHz alarm calls in the imminent paradigm (example on top) vs. shorter, more complex high frequency calls in the remote paradigm (example on bottom). (**I**) In both models, a fraction of central amygdala (CeA) cells are activated, as measured with c-Fos immunohistochemistry (white: home cage control, red: imminent, blue: remote). Number of animals, C-F: imminent threat, n=10, control group, n=10; remote threat, n=10, control group, n=8; G,H: imminent threat, n=9, control group, n=9; remote threat, n=5, control group, n=4; I: imminent threat, n=7, control group, n=10; remote threat, n=6, control group, n=8, home cage, n=6; Bars indicate mean ± standard error of the mean (SEM), * p < 0.05, ** p < 0.01, *** p < 0.001.

### Socially triggered passive and active defensive strategies are mediated by distinct neuronal circuits in the central amygdala

The behaviors displayed by observers (freezing, flight, and rearing) are known to be controlled by the CeA in the context of first-hand threat exposure^12^. We decided to investigate if similar circuitry controls reactions which are triggered socially. Staining for c-Fos, an immediate-early gene product used as a proxy for neuronal activation^14^, revealed that both imminent and remote paradigms activated a significant proportion of CeA neurons in comparison with home cage control animals (Fig. 1I; imminent threat: U=4, p=0.0140, remote threat: t=4.738, df=10, p=0.0008). To selectively manipulate the activity of cells involved in each paradigm^15^, we developed viral vectors carrying channelrhodopsin (ChR2) or halorhodopsin (eNpHR3.0) under the *c-fos* gene promoter (Fig. S1-4). This approach allowed us to tag populations of cells activated by imminent or remote paradigms and to stimulate or inhibit them 24 hours later in a different behavioral test (Fig. 2A,B). To investigate the effect of optogenetic manipulation on defensive responses, we designed an exploration test in which rats could freely explore novel, unfamiliar surroundings or avoid them by hiding in a dark shelter (Fig. 2C). Risk assessment, i.e., defensive behavior displayed towards aversive stimuli, was measured here as a stretched attend posture^16^ in the brightly lit corner of the test area.

**Fig. 2.**
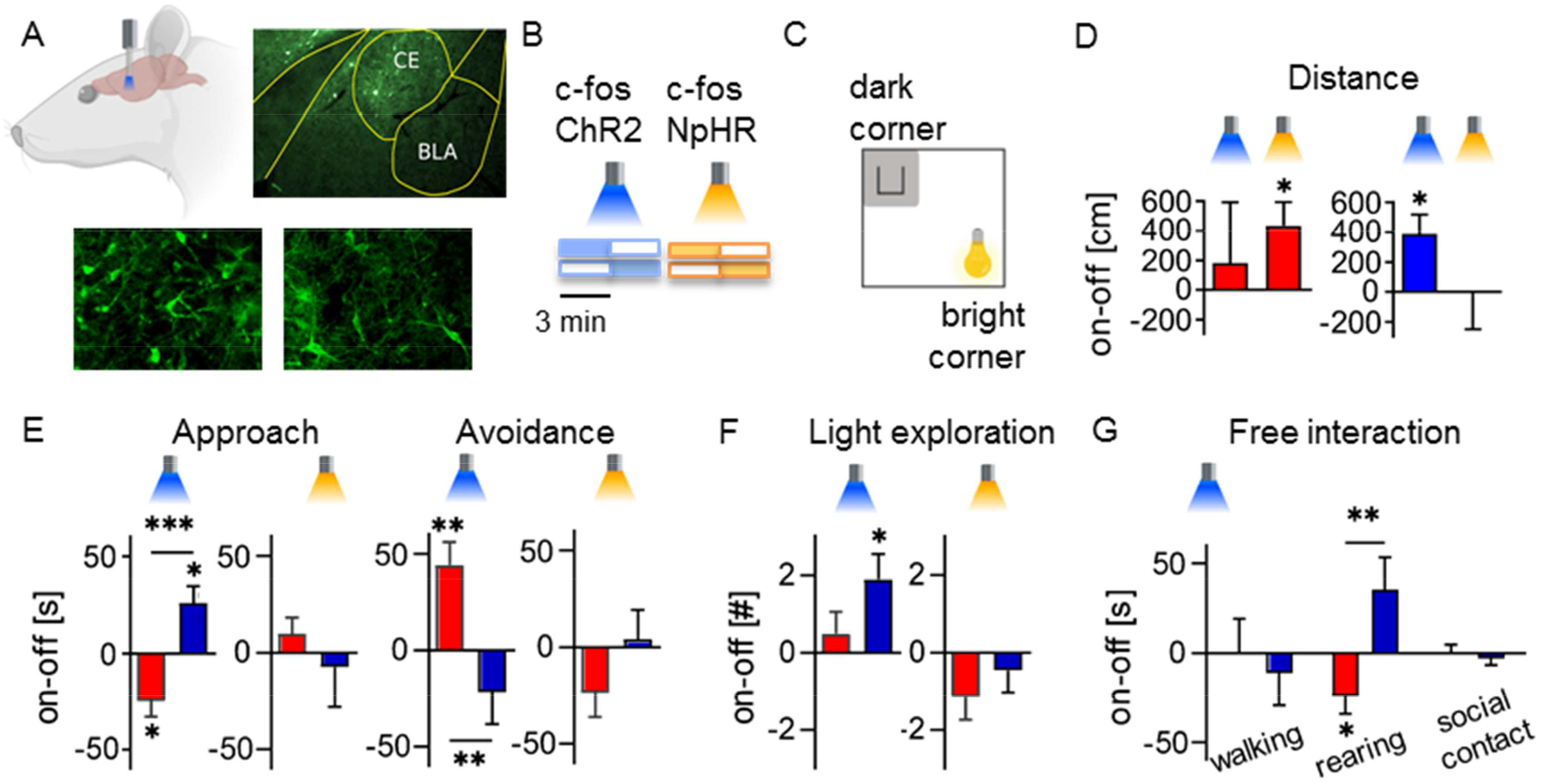
Optogenetic manipulations reveal different functions of subpopulations activated by imminent vs. remote threat paradigms. (**A**) To manipulate the responses of cells activated by each paradigm, we expressed ChR2 (bottom left) or eNpHR3.0 (bottom right) in a c-Fos dependent manner in observers’ CeA (upper right). (**B**) Laser stimulation was provided in 3-min long, counter-balanced ON-OFF periods (pulsed in the case of ChR2). (**C**) Schematic representation of open field used for measuring approach (exploration) and avoidance behaviors. (**D**) Rats covered a longer distance after stimulation of „remote” or inhibition of „immediate” subpopulation. (**E**) Stimulation/inhibition reveals opposite functions of the two subpopulations in controlling approach and avoidance. (**F**) Rats perform more episodes of light exploration after stimulation of the „remote” subpopulation. (**G**) Activation of each group of cells influences exploratory, but not social behaviors. Number of animals, exploration test: ChR2-imminent threat, n=8, NpHR-imminent threat, n=7, ChR2-remote threat, n=9, NpHR- remote threat, n=10; social interaction test: remote threat, n=8, imminent threat, n=8. Bars indicate mean ± standard error of the mean (SEM), * p < 0.05, ** p < 0.01, *** p < 0.001.

Optical stimulation or inhibition of neurons activated by either the imminent or remote paradigm affected exploration in a way consistent with the behavioral model. Rats generally moved more when ‘remote’ neurons were stimulated (t=2.955, df=8, p=0.0183) or ‘imminent’ neurons inhibited (Fig. 2D; t=2.656, df=6, p=0.0377). A more detailed analysis of time spent in different parts of the experimental area showed that activation of the ‘imminent’ neurons decreased exploration (defined as locomotion or rearing in the middle and lighted part of the arena; t=3.044, df=7, p=0.0187) and increased avoidance (measured as time spent in the shelter; t=3.636, df=7, p=0.0083). Stimulation of the ‘remote’ neurons resulted in increased exploration (Fig. 2E; t=2.930, df=8, p=0.0190; between-group comparisons for ChR2 exploration: t=4.179, df=15, p=0.0008, and for ChR2 avoidance: U=8, p=0.0053). In particular, during stimulation of ‘remote’ cells, rats performed more approaches to the brightly lit part of the arena (Fig. 2F; t=2.884, df=8, p=0.0204). Because these episodes were short (Fig. S5C), often accompanied by stretch-attend postures and followed by escapes (cf. Supplementary Movies)^16^, we interpret them as risk assessment. Changes in the exploration of the environment were not observed when neurons activated by contact with a partner that was not emotionally aroused were stimulated or inhibited (Fig. S5).

To test whether activation of the ‘imminent’ and ‘remote’ neurons affects social interactions, we stimulated these neurons during interaction with a partner. The results show that, though activated by social signals, the ‘imminent’ and ‘remote’ neurons modulate exploration of the environment rather than social interaction (Fig. 2G, S6; group x behavioral measure interaction: F=(2,28)=4.649, p=0.0181, post-hoc, rearing: p=0.0039; remote threat, rearing: t=2.379 df=7, p=0.0490). Since the area used for exploration testing was too big to reliably detect freezing, we decided to stimulate ‘imminent’ neurons in the context of a small, confined cage. Such manipulation did increase freezing, reproducing behavior from the imminent threat paradigm. This suggests that the activation of ‘imminent’ neurons leads to different defensive reactions depending on the context (Fig. S7). In summary, manipulation of activity-defined cell ensembles in the CeA revealed the existence of functionally defined ‘imminent’ and ‘remote’ subnetworks, which modulate defensive behavior in a context-specific manner. Activation of the ‘imminent’ network generally reduces movement while activation of the ‘remote’ network increases risk assessment behavior.

### Neuronal circuits activated by imminent vs. remote danger differ in molecular markers and patterns of connectivity

To test if ‘remote’ and ‘imminent’ cells can be identified based on molecular markers, we performed double immunostainings for c-Fos and corticotropin-releasing factor (CRF). The latter was recently found in fear-promoting circuits ^17^, in particular in CeA neurons that controlled active, but not passive, fear responses in individually-tested mice ^3^. Consistent with this data, the proportion of c-Fos-expressing (active) neurons containing CRF increased in the remote paradigm (CeL: t=2.466 df=6, p=0.0487; CeM: t=2.502 df=6, p=0.0464) but not in the imminent paradigm (compared to respective control groups; Fig. 3A). In contrast, the proportion of active PKC-δ neurons, recently shown to control the level of freezing^2^, was decreased in the remote paradigm (CeL: t=9.345 df=8, p<0.0001; CeM: t=2.867 df=8, p=0.0209) but not in the imminent paradigm (Fig. 3B). These results confirm that the populations of CeA cells activated in the imminent and remote paradigms are different.

**Fig. 3.**
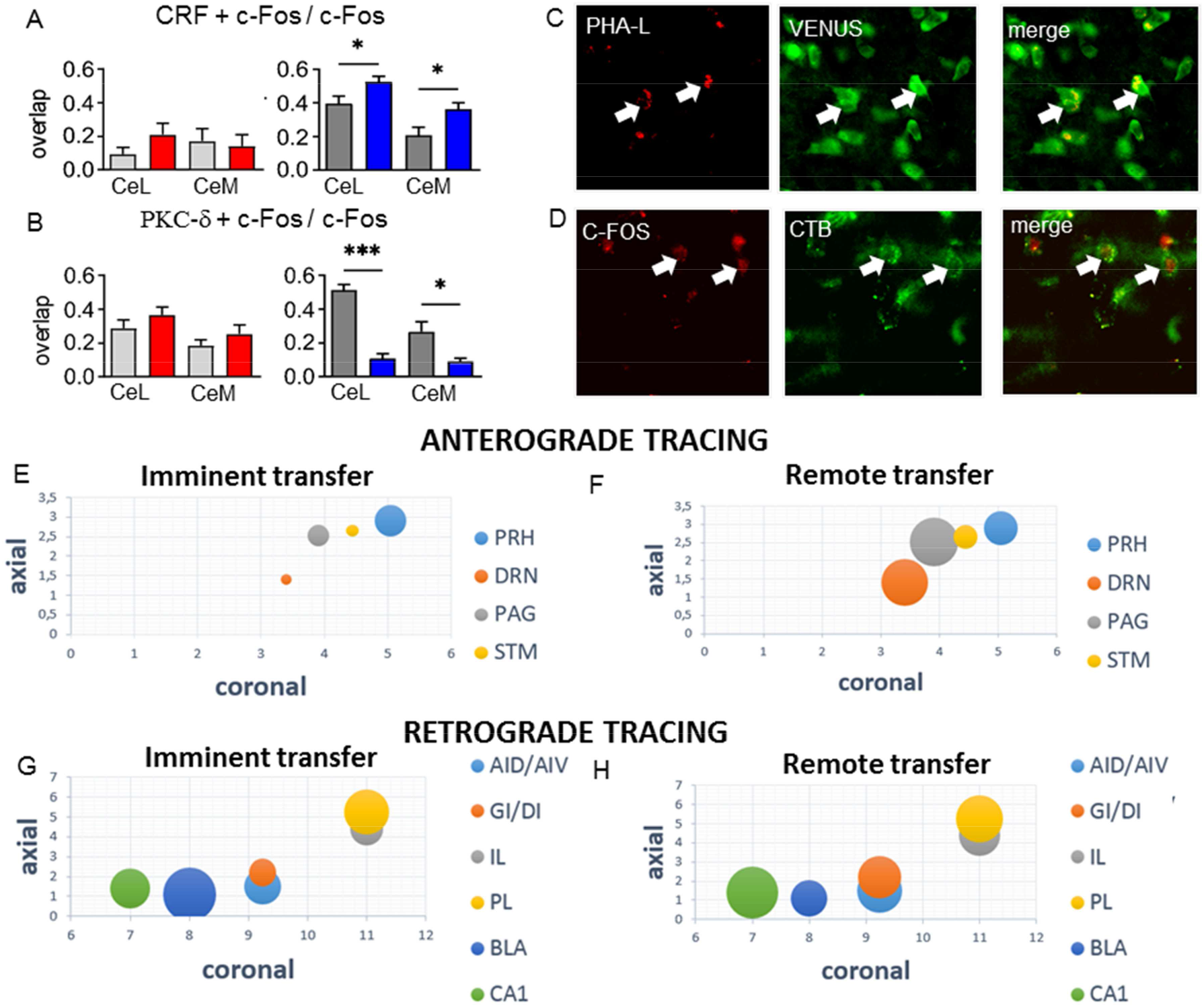
Cells activated by imminent vs. remote threat paradigm have different molecular markers and connectivity patterns. (**A**) Cells activated by the remote (blue), but not the imminent (red) paradigm express CRF (gray: control groups). (**B**) Cells activated by the remote, but not the imminent paradigm do not express PKC-δ. (**C**) Sample of Venus-positive neurons receiving projections from the CeA (anterograde tracing). (**D**) Sample of c-Fos-positive neurons sending projections to the CeA (retrograde tracing). (**E/F**) The number of activated neurons that receive projections from the CeA is proportional to the diameter of the area of the bubble, x and y values represent coronal and axial stereotaxic coordinates, respectively [Paxinos] (**G/H**) Percentage of activated neurons that send projections to the CeA is proportional to the diameter of the area of the bubble, x and y values represent coronal and axial stereotaxic coordinates, respectively [Paxinos]; CeL - lateral part of the CeA, CeM - medial part of the CeA, AID/AIV - agranular insular cortex dorsal/ventral, GI/DI - dysgranular insular cortex/granular insular cortex, IL - infralimbic cortex, PL - prelimbic cortex, BLA - basolateral amygdala, CA1 - field CA1 of the hippocampus PRH - perirhinal cortex, DRN - dorsal raphe nucleus, PAG - periaqueductal gray, STM - bed nucleus of the stria terminalis. Bars indicate mean ± standard error of the mean (SEM), * p < 0.05, *** p < 0.001.

To evaluate if ‘imminent’ and ‘remote’ neurons project to different downstream targets, we performed functional tracing using c-fos-PSD-95:Venus transgenic rats (Fig. 3C,D, Fig. S8). The PSD-95:Venus construct encodes fusion of the PSD-95 protein (a major component of postsynaptic densities) and the Venus reporter protein, followed by the Arc 3’UTR - dendrite localization sequence. This construct allows the dendrites and synapses of activated neurons to be visualized with fluorescent tags^18^. Combined with injection of anterograde tracer, PHA-L, we were able to visualize how many neurons were activated in multiple brain regions and which of them were receiving projections from the CeA. We show that the pattern of activation of the structures downstream from the CeA differs between the imminent and remote models. The number of active neurons which received projections from the CeA was typically higher in the remote paradigm, especially in structures such as the dorsal raphe nucleus, bed nucleus of the stria terminalis, periaqueductal gray, and perirhinal cortex (Fig. 3E,F, Table S1). The active CeA inputs also differed between the imminent and remote threat paradigms, with stronger activation of the basolateral amygdala projections in the imminent threat model and the infralimbic and insular cortex projections in the remote threat model (Fig. 3G,H, Table S2).

## Discussion

We discovered that animals are able to socially transfer specific information about distance to threat and that passive and active responses to social signals about threats are controlled by two distinct populations of CeA neurons. The first population is activated by signals about imminent danger and triggers passive defensive reactions: freezing and immobility. It is PKC-δ positive and most probably receives important input from basolateral amygdala. The second population, activated by information about remote threats, controls active risk assessment. It is PKC-δ negative and releases CRF. Its activity is most probably driven, among other structures, by the insula and infralimbic cortex.

These findings are important for understanding the mechanisms of emotional contagion for several reasons. Firstly, emotional contagion is typically studied using simplified paradigms, in which freezing of one rodent triggers freezing of the other in an apparently automatic manner. Here we show that animals are able to use social information much more flexibly when exploring a novel environment. Using socially transferred information about proximity of threat and habitat features, they choose the most appropriate defensive strategies. Secondly, our results are consistent with literature on single-subject defensive reactions. It is known that within mouse CeA, PKC-δ expressing circuits control freezing^2^, whereas CRF is implicated in active defensive responses^3^. The data support the hypothesis that social and non-social information about threat is processed, at least partially, through the same systems in the CeA^3,19^. This is an important finding because it has long been hypothesized that engaging the same circuits in first-hand and social experience provides a mechanism for complex social cognition, i.e. perspective taking^19^. Finding such candidate circuits in rodents provides a unique chance for verifying this hypothesis causally.

Thirdly, we studied animals which were experimentally naïve - that is, they were never subjected to any aversive manipulation before the imminent/remote paradigms. In many emotional contagion experiments it is common practice to pre-expose observers to foot-shocks in order to augment their reaction to social cues^20,21^. Our data proves that social information can elicit robust and flexible behavioral responses without any prior training.

According to a well-established theoretical model^22^, emotional contagion is a prerequisite for higher order social skills, including empathy. However, if one reduces this form of communication to pure mimicry, it seems unclear why it should support perspective taking or consolation behaviors^23,24^. We demonstrate that even naïve rodents are able to use basic socio-emotional cues to flexibly regulate their behavior. This suggests that emotional contagion can be a foundation for more sophisticated forms of social interaction.

## Materials and methods

### Experimental Design

Our study aimed to (i) systematically compare behavioral changes and ultrasonic communication (USVs) in rats exposed to social cues indicating either imminent or remote danger, (ii) determine the role of neuronal populations in the central amygdala (CeA) activated by social cues indicating either imminent or remote danger, (iii) characterize functional anatomy of neuronal circuits activated by social cues indicating either imminent or remote danger. Accordingly, we analyzed behavior and USVs in rats tested in imminent and remote threat models, tested the effects of optogenetic stimulation and inhibition of the CeA circuits activated by social cues indicating imminent vs. remote danger during exploration of a novel environment and social interaction, and identified molecular markers and connectivity of the activated circuits. In all experiments, rats were randomly assigned to experimental groups. In the experiments aiming at behavioral assessment (i) the groups consisted of 8-10 rats and all animals were included in the analysis. For the USV analyses, all data successfully recorded were used for the analyses. For c-Fos expression analysis, groups of 6-10 rats were used and the expression was analyzed (blindly) for all tested animals. In the optogenetic experiments sample sizes were established prospectively as 6-10 rats per group and in tracing experiments as 3-4 rats per group. We performed two replications of each optogenetic experiment, with ON and OFF periods order counter-balanced. The rats with misplaced viral vector/tracer injections or optic cannulas were eliminated from the analyses. The behavioral experiments which were manually scored were analyzed blindly by two trained observers.

### Animals

Experimental subjects were adult, experimentally naïve male Wistar rats (250–300 g at the beginning of the experiment), supplied by the Center of Experimental Medicine in Bialystok, Poland. The animals were randomly paired and housed together in standard home cages (43.0 × 25.0 × 18.5 cm) under a 12/12 light–dark cycle, with food and water provided *ad libitum*. The rats were habituated to the experimenter’s hand for 14 days preceding the experiment. Subjects underwent habituation to the transport, experimental room, and separation 3 days prior to the experiment.

For axonal tracing experiments, subjects were male c-Fos-PSD95Venus-Arc rats (300–400 g at the beginning of the experiment) bred at the Nencki Institute Animal House or Faculty of Biology Animal House (University of Warsaw). Rats were housed in pairs in standard home cages (43.0 × 25.0 × 18.5 cm) under a 12/12 light–dark cycle, with food and water provided *ad libitum*. The rats were habituated to the experimenter’s hand for 14 days preceding the experiment and to the transport, experimental room, and separation 3 days before the experimental day.

### Generation and genotypying of c-Fos-PSD95Venus-Arc Rats

All experiments were carried out in accordance with the Polish Act on Animal Welfare, after obtaining specific permission from the First Warsaw Ethical Committee on Animal Research.

### Viral vectors

#### Generation of c-Fos-ChR2/c-Fos-NpHR constructs and AAV packaging

To generate c-Fos-ChR2(H134R)-EYFP and c-Fos-NpHR-EYFP constructs (Fig. S1), we amplified a 1648 bp c-*fos* promoter fragment by PCR using Phusion High-Fidelity DNA Polymerase (ThermoFisher Scientific) from a previously described plasmid c-Fos-PSD95:Venus-Arc. The amplified *c-fos* sequence was inserted using 5’MluI and 3’KpnI restriction sites to substitute the synapsin 1 promoter in the pAAV-syn1-ChR2(H134R)-EYFP plasmid or to substitute the Thy1 promoter in the pAAV-thy1-NpHR-EYFP plasmid (both gifts from Karl Deisseroth). Cloning procedures were done in One Shot Stbl3 (ThermoFisher Scientific) competent cells to minimize recombination events. The resulting pAAV-c-Fos-ChR2(H134R)-EYFP and pAAV-c-fos-NpHR-EYFP plasmids were verified by sequencing. Following isolation using EndoFree Maxi Prep (Qiagen), the pAAV-c-Fos-ChR2(H134R)-EYFP and pAAV-c-fos-NpHR-EYFP plasmids were used to generate recombinant AAV vectors of mosaic serotype 1/2. Vectors were purified from cell lysates and their genomes were titrated by quantitative PCR.

#### Validation of c-Fos-NpHR construct in vitro

For activity control of the c-Fos-NpHR construct, hippocampal neurons were transfected with pAAV-c-Fos-NpHR-EYFP at DIV7 and stimulated with BDNF (10 ng/μl), KCl (50 mM), or chemically-induced LTP (cLTP: 50 μM forskolin, 50 μM picrotoxin, rolipram 0.1 μM) for 24 h at DIV20. Stimulation was also done in neurons in which neuronal activity was silenced with a sodium channel blocker (TTX, 1 μM), AMPA receptor antagonist (CNQX, 40 μM), NMDA receptor antagonist (APV, 100 μM), and a calcium channel blocker (nimodipine, 5 μM). To analyze the time course of NpHR expression, KCl (50 mM) was added to pAAV-c-Fos-NpHR-EYFP-transfected hippocampal neurons at different time intervals (Fig S3). Neurons were lysed with sample buffer and equal amounts of protein samples were resolved by SDS-PAGE. Following separation, proteins were semi-dry transferred onto PVDF membranes and probed with primary antibodies: anti-GFP (#598 MBL), anti-c-fos (E-8, mAb, SantaCruz) and GAPDH (mAb, EMD Milipore), and appropriate HRP-conjugated secondary antibodies. Immune-reactions were visualized using ECL (Amersham) (Fig. S2).

### Surgical procedures

#### Anterograde/retrograde tracing

Rats received intracranial injections of anterograde axonal transport tracer PHA-L Alexa Fluor 647 conjugate (Invitrogen; Molecular Probes) into the CeA, bilaterally, 14 to 21 days before behavioral training. In the case of retrograde tracing, the CeA of the rats were injected with 200 nl of retrograde tracer cholera toxin B Alexa Fluor 488 conjugate (CTB, Life Technologies, C34775). All surgical instruments were sterilized before surgery. Rats were anesthetized with isoflurane (5% induction, 1% for maintenance) and were given a subcutaneous injection of an analgesic (Butorfanol, 1 mg/kg). Ocular lubricant was used to moisten the eyes and the scalp was shaved. After being placed into the stereotaxic apparatus (David Kopf Instruments), the scalp was disinfected with 70% (vol/vol) alcohol, incised, and retracted. Two small burr holes were drilled to allow for a 1 μL NanoFil syringe needle (*World Precision Instruments*) to be lowered into the desired part of the brain. The coordinates used for the CeA were anteroposterior (AP), −1.8; mediolateral (ML), ±03.8; dorsoventral (DV), −7.5 [Paxinos]. The tracer PHA-L (2.5% wt/vol, dissolved in 0.1 M sodium PBS, pH 7.4) was delivered into the CeA (500nL per hemisphere, 100nL/min) using UMP3 UltraMicroPump (*World Precision Instruments*) and the syringe remained in place for another 5 min to allow for the diffusion of the tracer. In the case of retrograde tracing, the tracer CTB (1.0 mg/mL, dissolved in 0.1 M sodium PBS, pH 7.4) was delivered to the CeA using the same parameters as in the anterograde protocol. After the injection, the incision was sutured, and the animals were administered an analgesic (Tolfedine; 4 mg/kg; s.c.) and an antibiotic Baytril (4 mg/kg; s.c). To avoid dehydration, the animals were given 1 mL of 0.95% NaCl/100 g of body weight by s.c. injection. The rats were kept on a heating pad until they recovered from anesthesia before returning to their home cages.

#### c-fos-ChR2/c-fos-NpHR3 viral vectors

The rats were injected with c-fos-ChR2 (250-400 nL/site) or c-fos-NpHRR3 (350 nL/site) viral vectors, using the same surgical procedures as described above (100 nL/min) using a UMP3 UltraMicroPump (*World Precision Instruments*). The only difference was that optic cannulas (200 μm in diameter) were implanted bilaterally 0.3 mm above the injections and afterwards secured with 2 skull screws and dental cement.

### Behavioral paradigms

#### Remote threat paradigm

In each pair of rats, one subject was marked as a demonstrator and the other as an observer. In the experimental group, the demonstrators were taken from their home cages for aversive conditioning (Pavlovian contextual fear conditioning). When the demonstrators were trained, their partners were kept in the home cages in a different, sound-attenuated room. Immediately after the training, the demonstrators were placed back in their home cages and allowed to interact with the observers for 12 min. The control group was composed of rats treated in the same way, except that the demonstrators were exposed to the experimental cage without any training. Cameras and USV microphones (*UltraSoundGate, Avisoft*) were located above the experimental area.

#### Pavlovian contextual fear conditioning

The demonstrators were subjected to Pavlovian contextual fear conditioning in a fear conditioning chamber (*MedAssociates)*. The training consisted of a 2-min adaptation period and 9 footshocks lasting 1 sec and having 1 mA intensity, which were applied with interstimulus intervals of 59 sec. The animals were removed from the experimental cage 1 min after the last footshock was applied.

#### Imminent threat paradigm

Each pair of animals was retrieved from the animal facility in their home cage. The procedure was performed in a shuttle box (62.0 × 18.0 × 29.0 cm) that consisted of two identical opaque dark acrylic compartments separated by a perforated metal wall allowing the animals to see, hear, and smell each other. The cage was covered by a movable transparent acrylic ceiling. After 2 min habituation, the demonstrator received 10 foot-shocks (1 s, 1 mA, ITI 59 s) provided by a commercial stimulator (*MedAssociates*). In the control group, the foot-shocks were not provided. Because both rats in the control groups underwent identical treatment, they were both considered ‘observers’ and used as such for further procedures to reduce the total number of animals. The only exception was the social interaction experiment, in which only one animal from a pair could be stimulated optogenetically and the other one was considered the ‘demonstrator’ (see below). The behavior of the animals was video-recorded (the camera was located in front of the apparatus), and USV were collected with 1 to 4 microphones (*UltraSoundGate, Avisoft*). The chamber was cleaned with 70 % ethanol after testing each pair of animals. Behavior was recorded with WinTV software and each trial was recorded for later analysis using a camera positioned above the apparatus.

#### Exploration test

The exploration test apparatus was constructed of gray plywood and measured 1m x 1m with 40 cm walls. In one corner there was an uncovered rectangular shelter whose walls were constructed of plywood (10 cm × 15 cm). The opposite corner was illuminated with intense light forming a bright sphere contrasting with the remaining part of the arena (635 Lux). The apparatus was located in a sound-attenuating, dimly lit room.

Exploration test: approximately 24 h after the remote/imminent threat paradigm, the observers were placed in the shelter area of the apparatus. The laser stimulation (for details, see below) was provided in 3 min-long cycles (ON or OFF, order counter-balanced), and a total block of testing lasted 12 min. The arena was cleaned between each rat using 70 % ethyl alcohol. Behavior was recorded with WinTV software using a camera positioned above the apparatus. The analysis was done offline.

#### Social interaction

The social interaction test was performed approximately 24 h after the remote/imminent threat paradigm. The demonstrator rat was placed in a single cage with fresh bedding for 15 min in a separate room. Immediately after separation, the demonstrator was placed back in the home cage with an observer. The laser stimulation of the observers (for details, see below) was provided in 3 min-long cycles (ON or OFF, order counter-balanced), and a total block of testing lasted 12 min.

#### Home cage control

Rats were randomly paired and housed together in standard home cages. The rats were habituated to the experimenter’s hand and underwent the same viral vector surgeries as animals in the remaining groups, except on the final day they were not subjected to any behavioral stimulation.

#### Behavioral data analysis

The data from behavioral experiments was manually scored by trained observers, with frame-to-frame temporal resolution using BehaView Software (P. Boguszewski, http://www.pmbogusz.net/?a=behaview). The ethogram included: 1) for the remote threat paradigm and the social interaction test: rearing, locomotor exploration, contact with the partner (nose to butt, nose to head, following, crawling under, allo-grooming), self-grooming, evading partner, wrestling, prosocial (being groomed without protest), digging, quiescence; 2) for the imminent threat paradigm: rearing, locomotor exploration, freezing, social contact (i.e., touching partition between chambers with nose); 3) for the open-field test: avoidance (being in the dark shelter), rearing, locomotor exploration, running away from light, light exploration, and quiescence. Exploration test experiments were additionally analyzed with EthoVision XT9 (*Noldus*) to extract locomotor trajectories. In the imminent threat model, freezing (defined as at least 1s of immobility) was scored automatically in videos using BehaActive software (P. Boguszewski). USV recordings were analyzed with RatRec software (Miron Kursa, Adam Hamed).

#### Optogenetics in vivo

Custom-made optic cannulas (200 um fiber, 0.39NA, *Thorlabs*) were prepared before the surgeries and glued into M3 metal joints. Blue (472 nm) or yellow (589 nm) laser light was provided by fiber-coupled lasers (*CNI*), split by an optical rotary joint (*Doric Lenses*, 0.22NA) and delivered through armored patch cords (*Doric Lenses*). Power was adjusted to obtain 10 mW at the cannula tips. The laser was triggered by an *ArduinoUno* microcontroller to provide 3-min long LIGHT ON/OFF periods (blue: 5ms pulses, 30 Hz; yellow: continuous stimulation).

### Immunohistochemistry

#### Preparation of the tissue

Two hours after the last behavioral test (imminent/remote threat paradigm), rats received a lethal dose of morbital (133.3 mg/ml sodium pentobarbital, 26.7 mg/ml pentobarbital), and were transcardially perfused with ice-cold 0.1 M PBS (pH 7.4, *Sigma*) and 4% (wt/vol) paraformaldehyde (POCh) in PBS (pH 7.4). The brains were removed and stored in the same fixative for 24 h at 4°C, and subsequently immersed in 30% (wt/vol) sucrose at 4°C. The brains were then slowly frozen and sectioned at 40 μm on a cryostat. Coronal brain sections containing the amygdala were collected for immunohistochemistry.

#### c-Fos + CRF immunohistochemistry

For double enzymatic staining, the free-floating sections were washed with PBS, incubated (10 min) with hydrogen peroxide 0.3 %, and then blocked with 10% (vol/vol) NGS in PBS for 1 h. Then sections were incubated (4°C, for 4 days) with anti-CRF chicken antibody (*ABCAM*) and anti-c-Fos rabbit antibody (*Millipore*), diluted in PBS (both 1:1000).

On the 5th day, the sections were rinsed in PBS with 0.3% Triton X-100 (PBST) and incubated at room temperature with biotinylated secondary anti-rabbit antibody made in goat (Vector) 1:1000 in PBS for 2 h, then rinsed several times with PBST. Subsequently, the sections were incubated with avidin biotin complex (Vector) 1:1000 in PBS (prepared 0.5 h before use) for 1 h. Following several rinses in PBST, sections were stained with 3′-diaminobenzidine tablets (*Sigma*) enhanced with NiCl and washed. After overnight washing in PBST at 4 °C, sections were incubated in avidin biotin complex (*Vector*) 1:500 in PBS (prepared 0.5 h before use) for 1 h. Following several washes in PBS, sections were incubated for 2 h at room temperature with biotinylated secondary anti-mouse antibody made in goat (*Vector*) 1:500 in PBS for 2 h, rinsed in PBS, and stained with VIP Substrate Purple Kit (*Vector, SK-4600*).

Subsequently, sections were transferred to microscope slides, air-dried, immersed in xylene for 30 sec, and mounted using Entellan mounting medium (*Merck*).

Double fluorescent staining was performed on free-floating sections. The sections were washed with PBS and blocked with 5% (vol/vol) normal goat serum in PBST for 1.5-h at room temperature. Subsequently, sections were incubated with anti-c-Fos rabbit antibody (Millipore) and anti-CRF chicken antibody (ABCAM), diluted with 3% NGS in PBST at room temperature for 24 h or for 72 h at 4 °C. The next day, sections were rinsed with PBST, before 2-h incubation at room temperature with a secondary antibodies conjugated to Alexa Fluor 488 made in chicken (1:500; Invitrogen) and Alexa 555 made in rabbit (Invitrogen, 1:500) or Alexa 594 made in rabbit (Invitrogen, 1:1000). After several washes, the sections were mounted onto glass slides, overlaid with the Fluoromount G Medium, and covered with a glass coverslip.

#### c-Fos + PKC δ immunohistochemistry

Immunofluorescence staining was performed on free-floating sections. The sections were washed with PBS and blocked with 5% (vol/vol) normal goat serum in PBST for 1.5-h at room temperature. Subsequently, sections were incubated with anti-c-Fos rabbit antibody (Millipore) and anti-PKC δ mouse antibody (BD Transduction Laboratories), diluted with 3% NGS in PBST at room temperature for 24 h or for 72 h at 4 °C. The next day, sections were rinsed with PBST, before 2-h incubation at room temperature with a secondary antibodies conjugated to Alexa Fluor 488 made in mouse (1:500; Invitrogen) and Alexa 555 made in rabbit (Invitrogen, 1:500) or Alexa 594 made in rabbit (Invitrogen, 1:1000). After several washes, the sections were mounted onto glass slides, overlaid with the Fluoromount G Medium, and covered with a glass coverslip.

#### GFP immunohistochemistry

GFP fluorescent staining was performed on free-floating sections. The sections were washed with PBS with 0.3% Triton X-100 (PBST), blocked with 10% (vol/vol) normal goat serum in PBST, and incubated overnight at 4°C with anti-GFP rabbit antibody (Invitrogen) diluted with 1% normal goat serum (NGS) in PBST. The next day, sections were rinsed with PBST before 1 h incubation at room temperature with a secondary antibody conjugated to Alexa Fluor 488 (1:500; Invitrogen). After several washes, the sections were mounted onto glass slides, overlaid with the Fluoromount G Medium, and covered with a glass coverslip.

#### c-Fos immunohistochemistry

Immunofluorescence staining for c-Fos was performed on free-floating sections. The sections were washed with PBS and blocked with 5% (vol/vol) normal goat serum in PBST for 1.5-h at room temperature. Subsequently, sections were incubated with anti-c-Fos rabbit antibody (*Millipore*) diluted with 3% NGS in PBST at room temperature for 24 h. The next day, sections were rinsed with PBST, before 2-h incubation at room temperature with a secondary antibody Alexa 555 made in rabbit (Invitrogen, 1:500). After several washes, the sections were mounted onto glass slides, overlaid with the Fluoromount G Medium, and covered with a glass coverslip.

#### Image Capture and Analysis

The double-labeling results were analyzed with the aid of a Nikon Eclipse Ni-U fluorescent microscope equipped with a QImaging QICAM Fast 1394 camera and an Ar laser producing light at 467 and 488 nm as well as a Kr laser for 568 nm. Two objectives (20× and 10×) were used to capture the samples with the aid of Image-Pro Plus 7.0.1.658 (Media Cybernetics) software. The images were then processed with ImageJ software by experimenters who were blind to the treatments during image acquisition as well as during cell counting. PHA-L images were merged with Venus-stained cell bodies and proximal dendrites to analyze for the presence of close appositions between PHA-L and Venus positive neurons in the particular structures. The ratio of the Venus neurons receiving projections to the whole number of Venus positive neurons was calculated. CTB positive cell bodies were merged with c-Fos positive images to examine whether c-Fos positive cell nuclei are localized in CTB positive neurons in selected structures. The ratio of the CTB positive cell bodies with c-Fos nuclei to all CTB positive neurons was calculated. Similarly, molecular markers PKC δ and CRF were quantified in ImageJ software. The ratio of PKC δ +/c-Fos + to c-Fos+ and CRF+/c-Fos+ to c-Fos+ in the CeA was calculated.

#### Statistical analysis

Data are presented as mean ± SEM. In the Remote vs. Imminent model behaviors (social contact, rearing, horizontal exploration, freezing), the density of c-Fos-positive neurons in the CeA, USV frequencies, as well as ratios of PKC-δ+ c-Fos / c-Fos and CRF + c-Fos / c-Fos results were analyzed by unpaired t tests or Mann-Whitney tests. In the exploration test, intensity of exploration of the environment, duration of active exploration/time spent in the shelter in the test area, between-group comparisons for ChR2 exploration and for time spent in the shelter, and number of explorations of the brightly lit area were analyzed by comparisons to theoretical 0 value with one sample t-tests, between-group comparisons with unpaired t tests, or with the Mann-Whitney test. The effect of the stimulation of the IT/RT neurons on walking, rearing, and social contact during social interaction was analyzed by ANOVA with repeated measures followed by Fisher’s LSD test. The nonparametric tests were used when data did not follow a normal distribution. All statistical analyses were completed using GraphPad Prism version 6 (GraphPad Software, Inc., San Diego, CA), p < 0.05 was considered significant. Illustrations were created using BioRender webpage.

## Author contributions

EKnapska designed research; KA, KK, KRS, KZS, TN, MW, KD, MW, DD performed research; EKublik, MB, TL, TJ, HM, WK, PMB contributed to the development of new reagents/analytic tools; KA, KK, KRS, KZS, KM, AH, MK, EKnapska analyzed data; and KK, KA, EKnapska wrote the paper.

## Data and materials availability

The datasets generated during and/or analyzed during the current study are available from the corresponding author on reasonable request.

## Other

We are thankful to Thomas G. Custer and Balazs Hangya for the critical reading of the manuscript.

