## Supplementary material for "More than a feeling: central amygdala mediates social transfer of information about proximity of danger"

**List of Supplementary Materials:**

**Figures S1-S8**

**Tables S1-S2**

**Movies S1-S6**

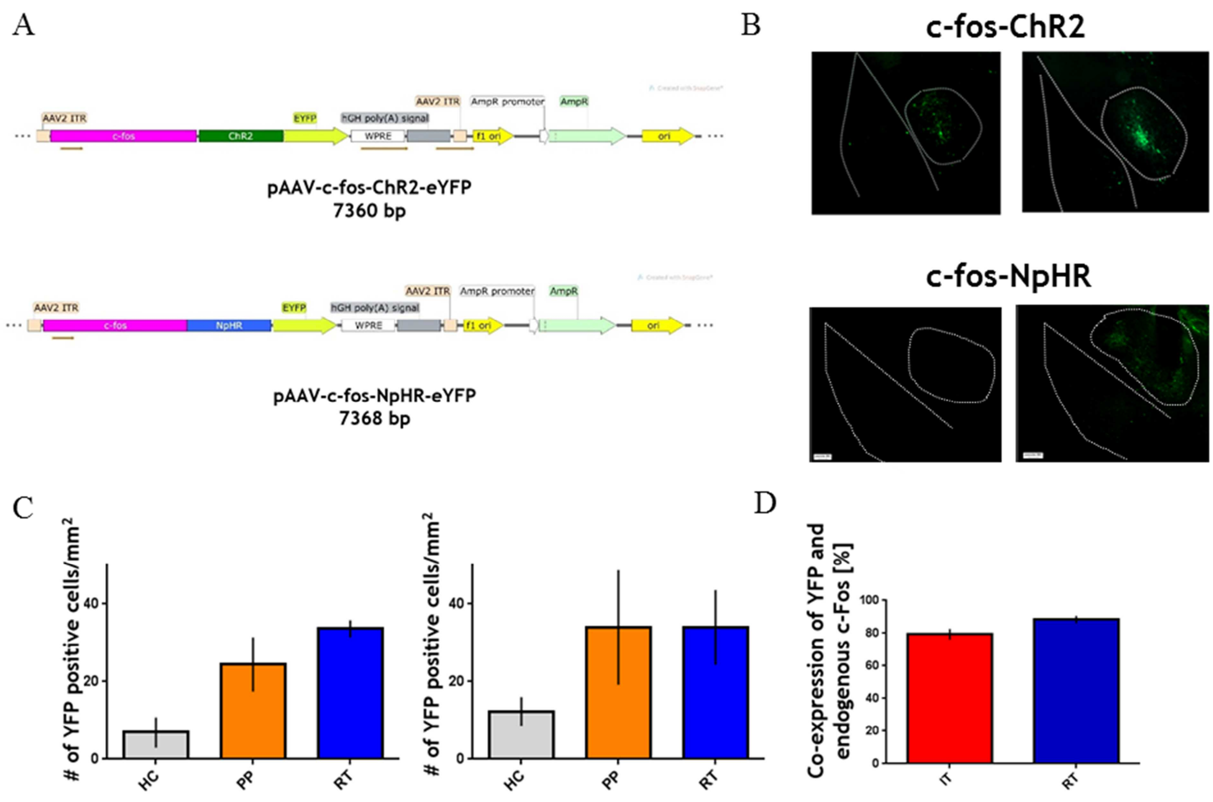


**Fig. S1. (A) Maps of the c-fos-ChR2 and c-fos-NpHR constructs. (B) Expression of fluorescent reporter protein in rats injected with c-fos-ChR2 or c-fos-NpHR constructs. For each case - left panel: non-stimulated (control) rat; right panel: behaviorally stimulated rat. (C) Expression of reporter protein in c-fos-ChR2 construct (left panel) or c-fos-NpHR construct (right panel) injected rats: home cage controls (H), after place preference paradigm (PP) and rats after remote threat paradigm (RT). (D) Co-expression level of the reporter protein and endogenous c-Fos in the CeA of stimulated rats (RT – remote threat paradigm, IT – imminent threat paradigm).**

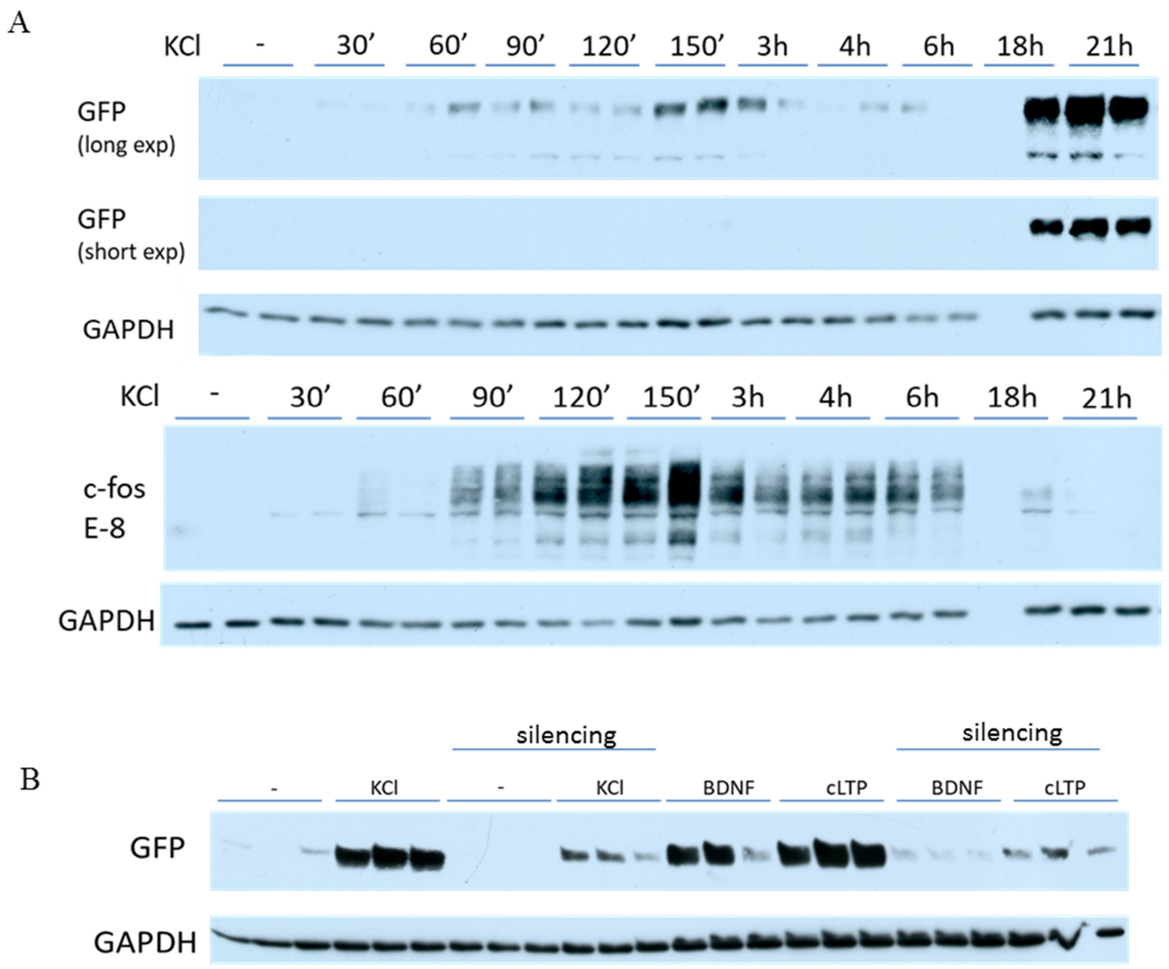


**Fig. S2. (A) Upper panel: western blot analysis of time course of NpHR-EYFP expression in hippocampal neurons after neuronal depolarization (KCl, 50 mM). Lower panel: time course of endogenous c-fos accumulation following neuronal depolarization (KCl, 50 uM). (B) Western blot analysis of NpHR-EYFP expression following BDNF, KCl and cLTP stimulation (20 h incubation) in the presence of TTX, CNQX, APV and nimodipine.**

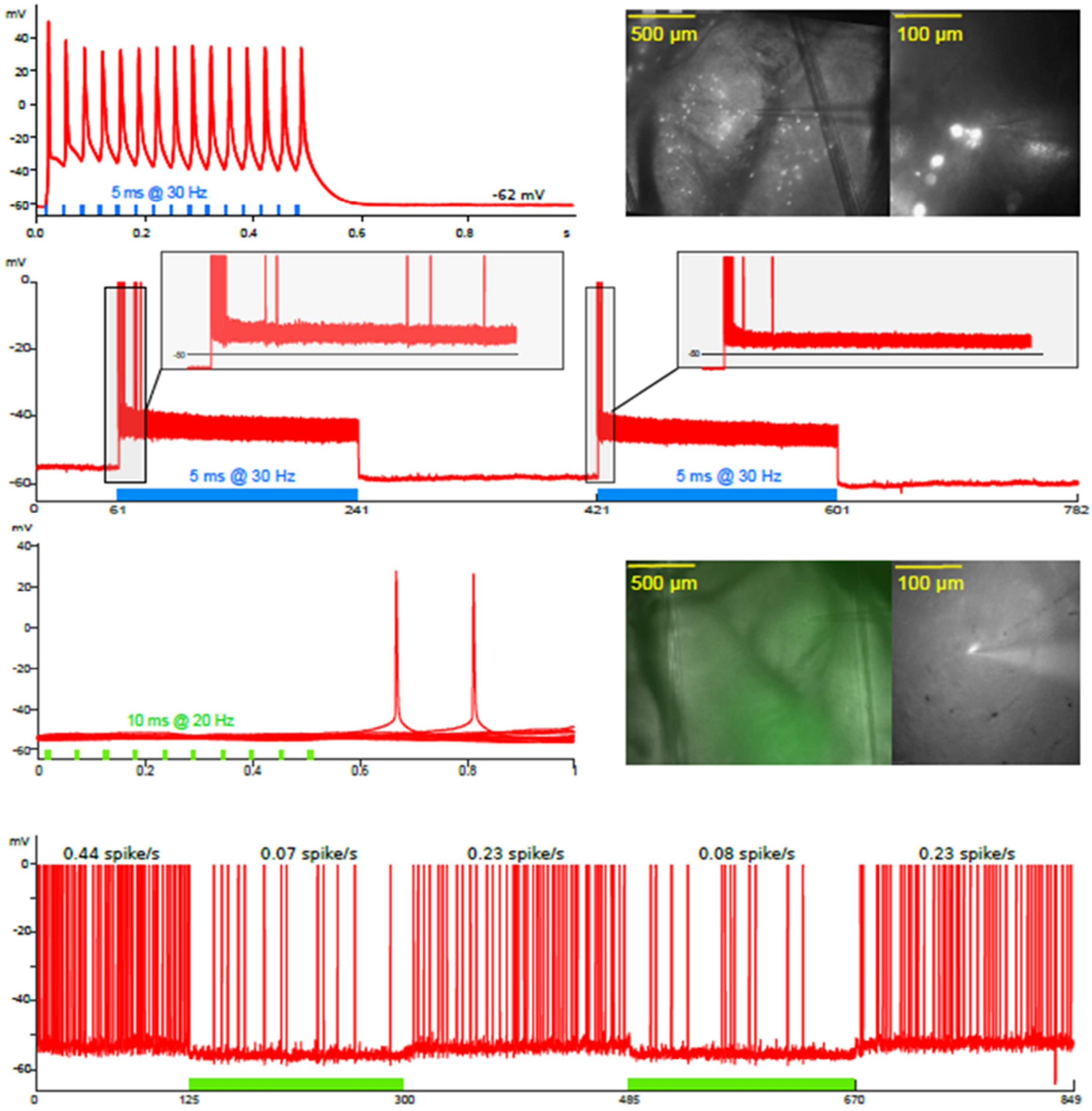


**Fig. S3. Whole-cell patch-clamp recordings from the neurons expressing the reporter protein. (A) Stimulation of neurons expressing c-fos-ChR2 construct with blue light (473 nm, in 5 ms pulses delivered at 30 Hz) resulted in cell activation. (B) Stimulation of neurons expressing c-fos-NpHR construct with amber light (591nm, in 10 ms pulses delivered at 20 Hz) resulted in cell inhibition.**

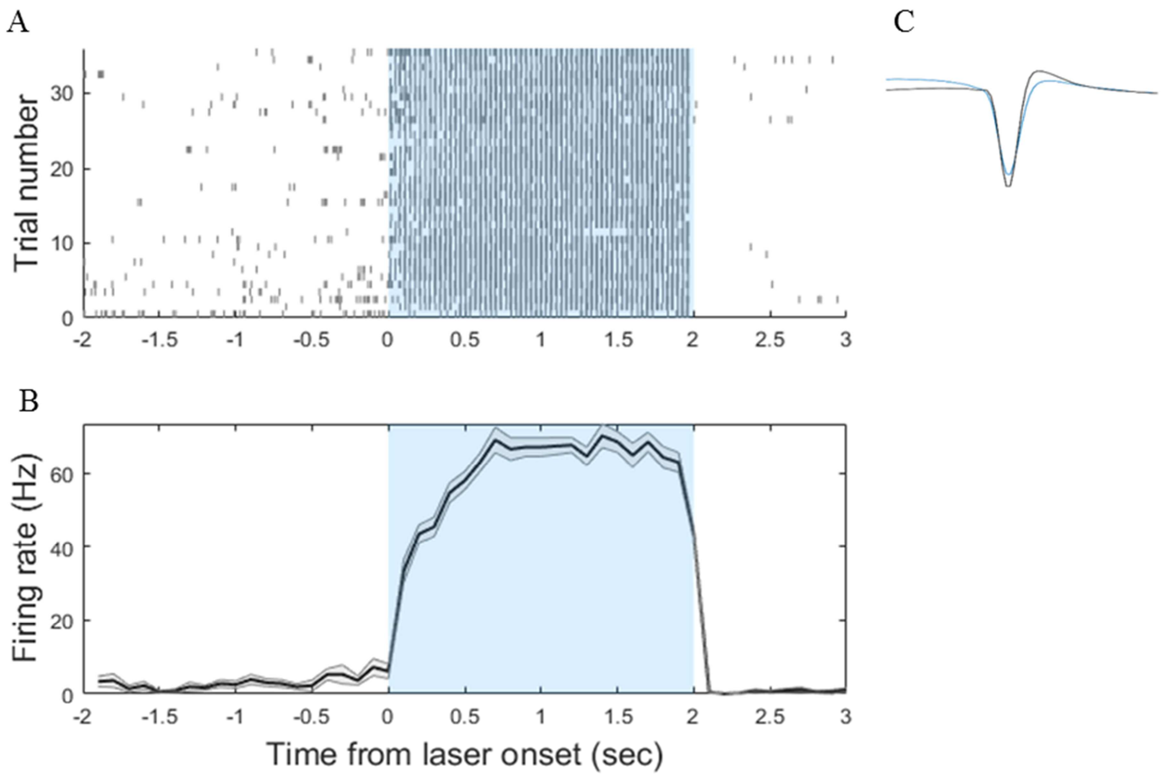


**Fig. S4. Effects of optogenetic stimulation on single unit recorded *in-vivo*. (A) Raster plot showing responses of exemplary neuron to laser stimulation (shaded in blue: 5 ms pulses, 30 Hz for 2 seconds, 10 mW). (B) Peri-event time histogram calculated from the same data; black line represents mean firing rate in each time bin (100 ms); area shaded in grey indicates standard errors of the mean. (C) Average waveform during light off (in black) and light on periods (in blue). The neuron was recorded from the dorsal hippocampus of a freely moving rat injected with the c-fos-ChR2 construct. During the same surgery, electrophysiological drive (8 tetrodes around 200 µm optic fiber) was implanted. Neurons were sorted with Kilosort2 and Phy; 5 ms-long periods during laser stimulation were excluded from the analysis due to contamination with artifacts.**

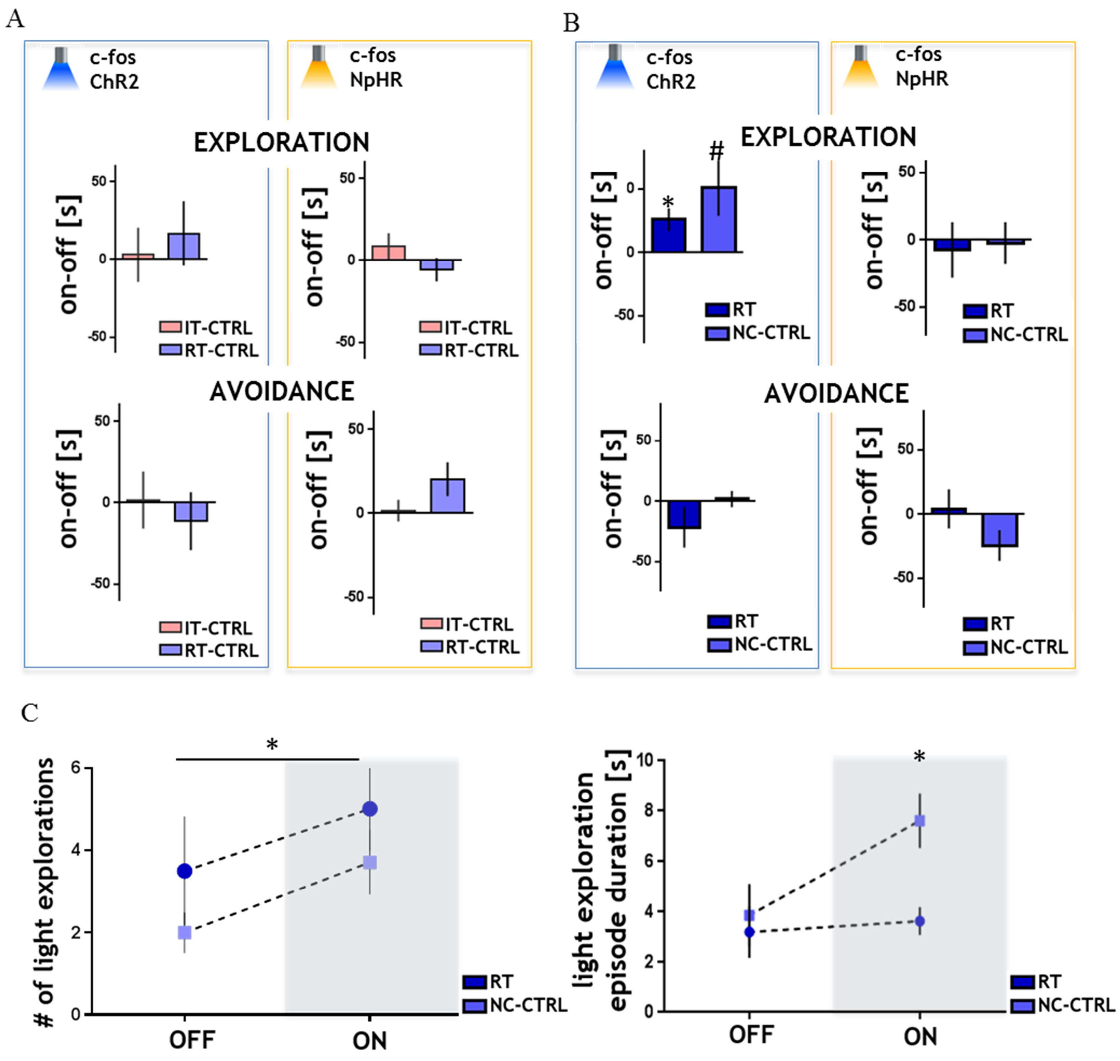


**Fig. S5. Effects of optogenetic stimulation and inhibition of CeA neurons activated by contact with a partner that was not emotionally aroused (NC-CTRL), assessed in the exploration test. c-fos targeted expression of ChR2 or NpHR was induced by interaction with a partner not subjected to fear conditioning yet exposed to the separated cage. (A) Analysis of time spent in different parts of the exploration test area - on active exploration and in the hiding box (avoidance) - showed no differences between the periods with and without stimulation or inhibition (ChR2-IT-ctrl, n=9, NpHR-IT-ctrl, n=10, ChR2-RT-ctrl, n=7, NpHR-RT-ctrl, n=9). (B) Interaction with partners exposed to a novel cage (NC-CTRL) showed a tendency toward increased exploration when stimulated optogenetically (t=2.319, df=6, p=0.0595, ChR2-NC-ctrl, n=7, NpHR- NC-CTRL, n=6). (C) Although the number of light explorations increases similarly in both groups [two-way ANOVA with repeated measures, the effect of on/off period: F (1, 11)=7.82, p=0.0174], the duration of light exploration increases in NC-CTRL group only [two-way ANOVA with repeated measures, the effect of group: F (1, 11)=5.862, p=0.0339 followed by Holm-Sidak's multiple comparisons test], indicating significantly lower anxiety level in the NT-CTRL animals. Bars indicate mean ± standard error of the mean (SEM), * p < 0.05, # p = 0.06 (comparisons to theoretical 0 value with one sample t-test or the Wilcoxon Signed Rank test, between-group comparisons with unpaired t test with Welch's correction or with the Mann-Whitney test).**

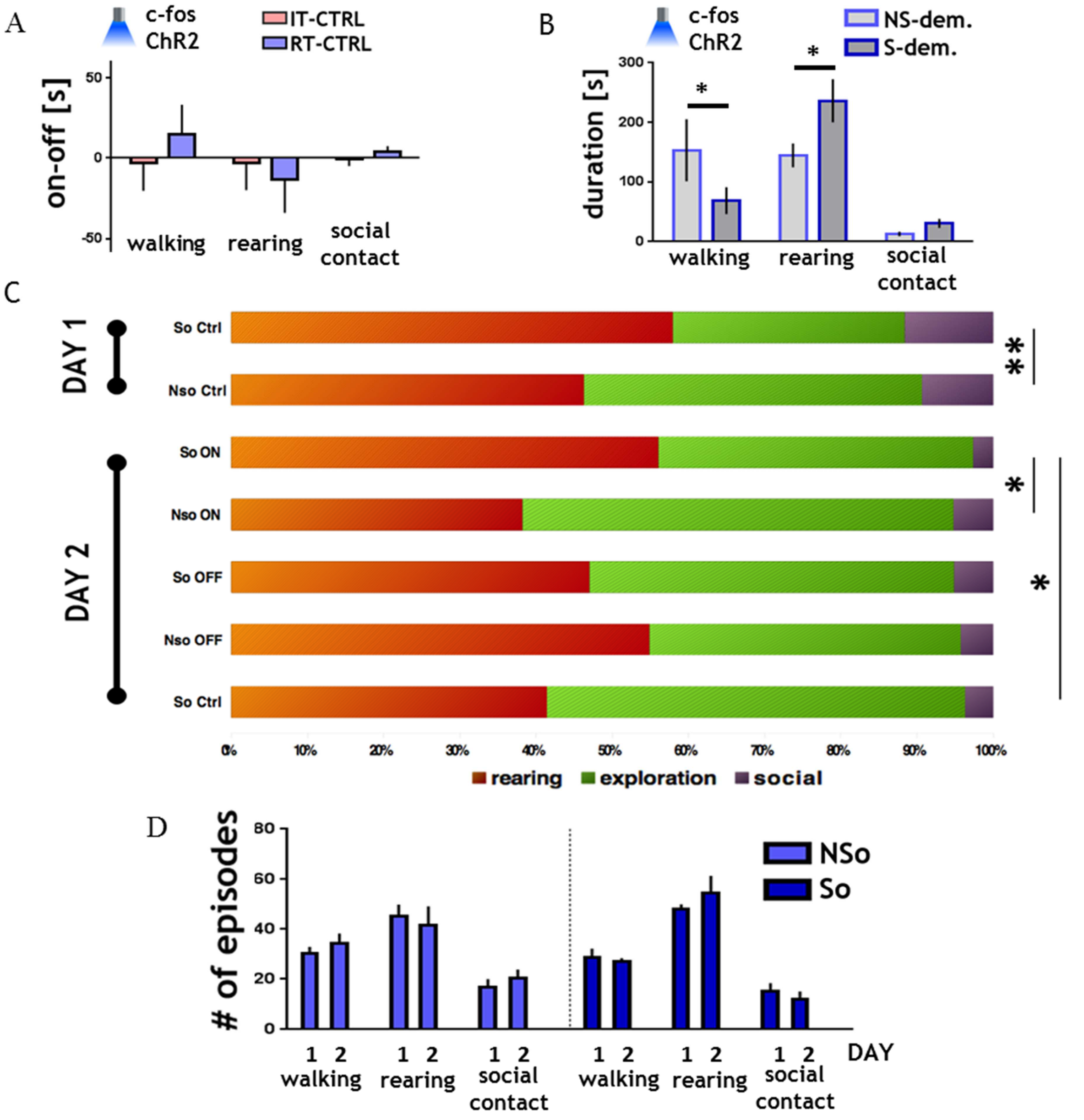


**Fig. S6. Effects of optogenetic stimulation of CeA neurons activated by contact with a partner that was not emotionally aroused on social interaction. Rats were injected with AAV-c-fos-ChR2-EGFP and implanted with optic fibers to the CeA. c-fos targeted expression of ChR2 was induced by interaction with a partner that was not subjected to**

**fear conditioning but placed in the same cage as rats in the IT or RT models. Effects of photostimulation on social interaction were tested. The graphs show a difference between 3-min periods with laser on and off. (A) Analysis of time spent on ambulation, rearing, and social contact showed no differences between the periods with and without stimulation. (B) When RT neurons in observers were optogenetically stimulated, as a secondary effect we also detected changes in behavior of demonstrators (who did not receive any optogenetic stimulation), NS_d – demonstrator interacting with observer from control group, S_d – demonstrator interacting with observer from RT group [ANOVA with repeated measures (behavioral measure), the group x behavioral measure: $F_{(2, 22)}=3.576$, $p=0.0452$ followed by Fisher LSD test]. (C) The comparison of behavioral changes during interaction with emotionally aroused partner (day1) with the effects of optogenetic stimulation (day 2) showed that photostimulation of the CeA RT neurons evoked behavioral changes similar to those observed during real interaction with an emotionally aroused partner, i.e., increased rearing; So - rats interacting with fear conditioned partners, n=8, NSo - rats interacting with non-stimulated partners, n=8, ON/OFF - light on/off; two-way ANOVA with repeated measures (session, behavioral measure), the group x session x behavioral measure interaction: $F_{(6, 126)}=4.0219$, $p=0.001015$ followed by planned comparisons; session day1, day2_ON, day2_OFF. Comparisons with behavior of rats that were not stimulated before interaction on the $2^{nd}$ day (So_ctrl, n=5) showed that the increased rearing is specific to optogenetic stimulation (factorial ANOVA: So_ON vs. So_ctrl, the group x behavioral measure interaction $F_{(2, 32)}= 3.3344$, $p=0.048374$). (D) Analysis of behavior during 10-min social interactions after short 10-min isolation periods. The interaction tests were repeated for 2 consecutive days. The data show that repeated social interactions evoked similar behavioral activity as the first test. Bars indicate mean ± standard error of the mean (SEM), * $p < 0.05$, ** $p < 0.01$ (comparisons to theoretical 0 value with one sample t-test or the Wilcoxon Signed Rank test, between-group comparisons with unpaired t test with Welch's correction or with the Mann-Whitney test; repeated social interaction with ANOVA with repeated measures (behavioral measure, session)).**

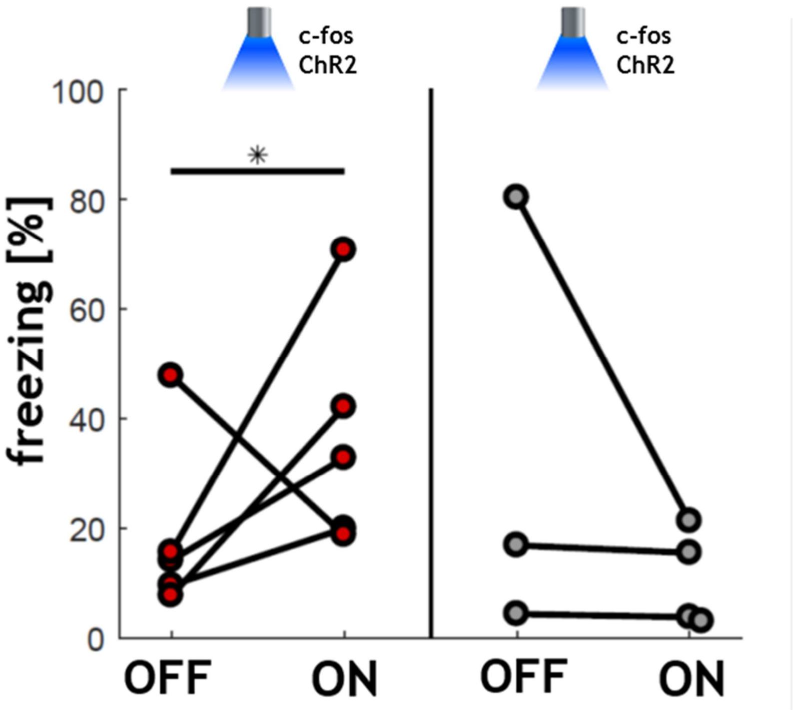


**Fig. S7. Stimulation of „immediate threat" neurons evoked freezing, when tested in confined cage (red data points: $p < 0.05$, one-tailed Wilcoxon test). There was no light effect in the control group (gray points; video from one OFF period was not recorded due to experimenter's error).**

**The rats (n = 5 exp, n = 4 ctrls), injected with c-fos-ChR2 construct underwent standard remote-threat (or control) paradigm. 24 hours later they were placed in the same experimental cage, and after 1 minute of habituation received 3min ON - 3 min OFF (counter-balanced) laser stimulation. Freezing was measured as described in the Methods section.**

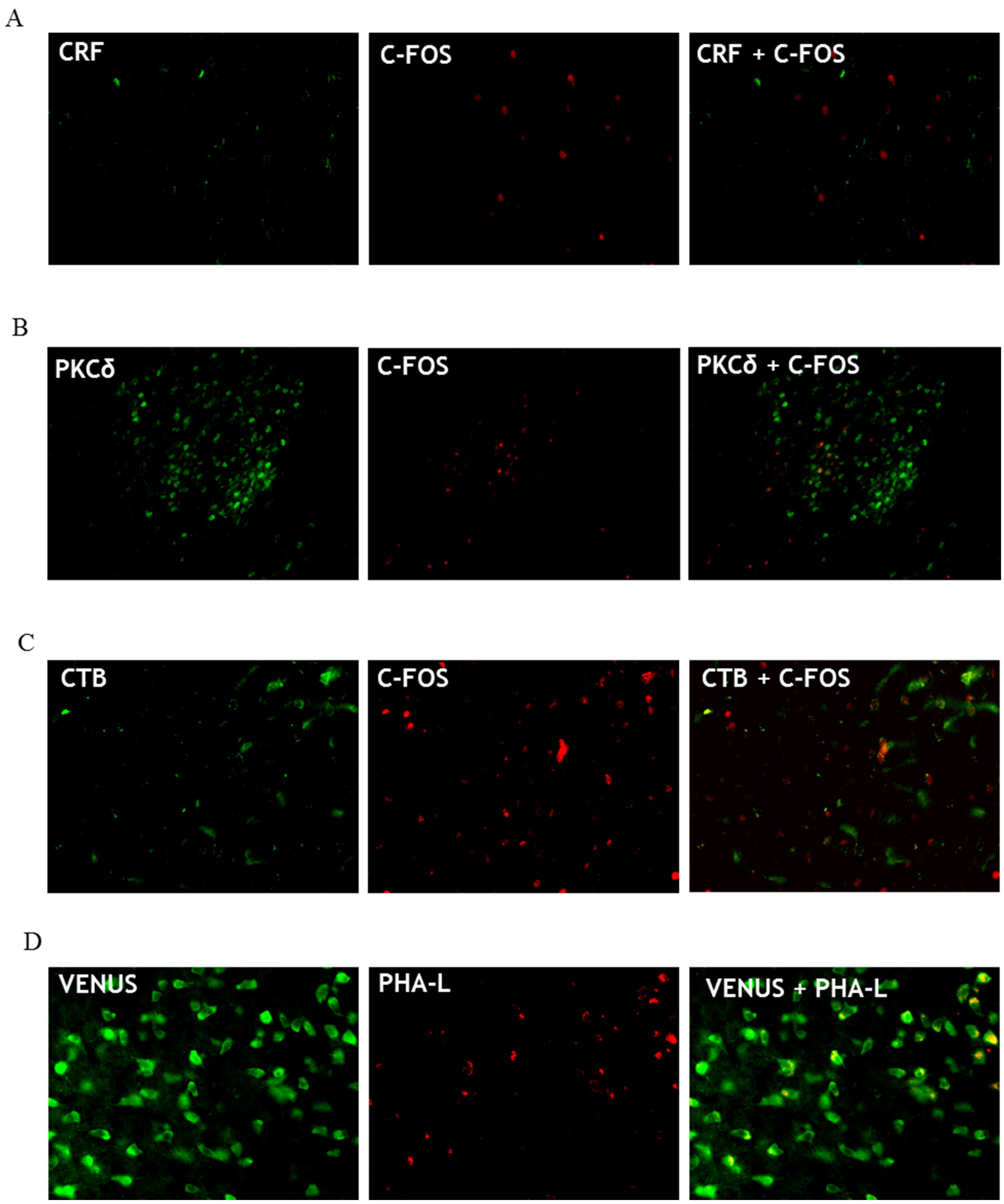


**Fig. S8. Representative examples of immunohistochemical stainings (see Methods for details). (A) Co-staining of CRF and c-Fos in CeA (B) Co-staining of PKCδ and c-Fos in CeA (C) Retrograde functional tracing: CTB-positive cells immunostained with anti-c-Fos antibody. (D) Anterograde functional tracing: sample immunostained with anti-GFP antibody and PHA-L anterograde tracer.**

**Tab. S1. Detailed results of the anterograde functional tracing experiment (summarized in Fig. 3).**

| brain structure | Imminent threat model | | Remote threat model | |
|---|---|---|---|---|
| | number of activated cells ± SEM | percent of activated cells receiving projections from the CeA ± SEM | number of activated cells ± SEM | percent of activated cells receiving projections from the CeA ± SEM |
| **BRAINSTEM** | | | | |
| dorsal raphe nucleus. dorsal part | 8.0 ± 1.0 | 78.7 ± 3.6 | 74.5 ± 1.8 | 100.0 ± 0.0 |
| dorsal raphe nucleus. ventral part | 4.3 ± 0.3 | 65.4 ± 9.6 | 124.3 ± 2.6 | 100.0 ± 0.0 |
| periaqueductal gray. ventrolateral part | 31.0 ± 4.4 | 77.2 ± 3.0 | 86.6 ± 2.0 | 100.0 ± 0.0 |
| periaqueductal gray. dorsolateral part | 12.8 ± 0.7 | 66.3 ± 6.9 | 50.3 ± 2.5 | 100.0 ± 0.0 |
| periaqueductal gray. lateral part | 15.8 ± 1.5 | 77.5 ± 4.3 | 50.3 ± 0.9 | 100.0 ± 0.0 |
| pedunculopontine tegmental nucleus | 11.9 ± 0.6 | 84.4 ± 6.5 | 30.8 ± 1.6 | 99.7 ± 0.3 |
| laterodorsal tegmental nucleus | 9.3 ± 1.2 | 76.6 ± 6.9 | 39.6 ± 7.4 | 94.1 ± 5.9 |
| substantia nigra | 34.7 ± 7.3 | 85.9 ± 1.1 | 135.1 ± 2.5 | 92.1 ± 4.5 |
| retrorubral field | 13.0 ± 2.1 | 77.8 ± 2.1 | 34.7 ± 3.6 | 92.2 ± 6.4 |
| ventral tegmental area | 6.3 ± 1.0 | 67.5 ± 1.6 | 22.3 ± 0.0 | 80.0 ± 0.0 |
| parabrachial nucleus | 4.1 ± 0.4 | 74.6 ± 5.0 | 34.3 ± 8.5 | 88.0 ± 12.0 |
| nucleus of the trigeminal nerve | 8.6 ± 1.5 | 98.1 ± 1.9 | 7.2 ± 0.2 | 100.0 ± 0.0 |
| **BASAL FOREBRAIN** | | | | |
| bed nucleus of the stria terminalis | 15.1 ± 5.8 | 78.3 ± 2.5 | 73.1 ± 21.4 | 79.1 ± 10.6 |
| globus pallidus | 42.8 ± 13.0 | 82.9 ± 3.7 | 30.4 ± 4.3 | 78.7 ± 0.6 |
| caudate putamen | 54.3 ± 7.3 | 80.9 ± 0.9 | 92.2 ± 8.5 | 78.7 ± 5.0 |
| substantia innominata | 13.1 ± 2.2 | 84.3 ± 3.5 | 45.0 ± 16.6 | 71.7 ± 10.5 |
| **HYPOTHALAMUS** | | | | |
| dorsal hypothalamic area | 5.7 ± 0.6 | 69.7 ± 5.8 | 27.8 ± 10.4 | 90.1 ± 6.8 |
| retrochiasmatic area | 38.7 ± 10.8 | 55.8 ± 10.7 | 56.5 ± 3.6 | 74.7 ± 13.8 |
| ventromedial hypothal nu | 15.9 ± 4.2 | 67.5 ± 3.2 | 41.1 ± 3.6 | 72.8 ± 13.2 |
| **CORTEX** | | | | |
| perirhinal cortex | 27.8 ± 4.6 | 80.0 ± 0.9 | 140.9 ± 24.4 | 80.5 ± 1.0 |

**Tab. S2. Detailed results of the retrograde functional tracing experiment (summarized in Fig. 3).**

| brain structure | Imminent threat model | | Remote threat model | |
|---|---|---|---|---|
| | number of retrogradely stained cells ± SEM | percent of cells sending projections to the CeA that are activated ± SEM | number of retrogradely stained cells ± SEM | percent of cells sending projections to the CeA that are activated ± SEM |
| **CORTEX** | | | | |
| Agranular insular cortex | 5,42 ± 1,06 | 13 ± 2 | 26 ± 2 | 4,25 ± 0,25 |
| Dysgranular insular cortex/granular insular cortex | 4±0,14 | 8 ± 0 | 23 ± 6 | 14,5 ± 5 |
| Infralimbic cortex | 6,33±1,02 | 11 ± 3.0 | 22 ± 8 | 11± 4 |
| Prelimbic cortex | 5,25 ±1,89 | 20 ± 6 | 28 ± 5 | 14,5 ± 5 |
| Cingulate cortex, area 1 | 6,08±1,47 | 37 ± 2 | 43 ± 22 | 11,5± 2,5 |
| **BASAL FOREBRAIN** | | | | |
| Basolateral amygdala | 36,25± 8,31 | 28 ± 6 | 18 ± 4,5 | 17 ± 2 |
| **HIPPOCAMPAL FORMATION** | | | | |
| Field CA1 of the hippocampus | 32 ± 2 | 15 ± 3 | 34 ± 3 | 13,5 ± 2 |

| | |
|---|---|
| **VIDEO 1: CTRL NS SOCIAL INTERACTION** | https://drive.google.com/a/nencki.edu.pl/file/d/18sRukV5VYbmVy04xg4jJSIr2D2eHGCDQ/view?usp=sharing |
| **VIDEO 2: CTRL NS EXPLORATION TEST** | https://drive.google.com/a/nencki.edu.pl/file/d/1ue61nGDM7wkzlatHTguzDEjgeaZIpNBa/view?usp=sharing |
| **VIDEO 3: IMMINENT SOCIAL INTERACTION** | https://drive.google.com/a/nencki.edu.pl/file/d/1N8YyQCZp66hRnDQGmP0KZJDot9nxH-FX/view?usp=sharing |
| **VIDEO 4: IMMINENT EXPLORATION TEST** | https://drive.google.com/a/nencki.edu.pl/file/d/1uEVcWaF9XVb_wXJt4gFD9tm6HY_venNY/view?usp=sharing |
| **VIDEO 5: REMOTE SOCIAL INTERACTION** | https://drive.google.com/a/nencki.edu.pl/file/d/1KiztNuurIUrusEUE282bFkec8srXUEs8/view?usp=sharing |
| **VIDEO 6: REMOTE EXPLORATION TEST** | https://drive.google.com/a/nencki.edu.pl/file/d/1PiYVEKyWfiHq_F_o8i9L-w81gB1obq5U/view?usp=sharing |
| **ALL** | https://drive.google.com/a/nencki.edu.pl/file/d/1wPkRrfwu5t6JQi9bOzM-O9r3GYHmSRyV/view?usp=sharing |
